# A second view on the evolution of flight in stick and leaf insects (Phasmatodea)

**DOI:** 10.1101/2021.10.12.464101

**Authors:** Sarah Bank, Sven Bradler

## Abstract

The re-evolution of complex characters is generally considered impossible, yet, studies of recent years have provided several examples of phenotypic reversals shown to violate Dollo’s law. Along these lines, the regain of wings in stick and leaf insects (Phasmatodea) was hypothesised to have occurred several times independently after an ancestral loss, a scenario controversially discussed among evolutionary biologists. Here, we revisit the recovery of wings by reconstructing a phylogeny based on a comprehensive taxon sample of over 500 representative phasmatodean species to infer the evolutionary history of wings. We additionally explored the presence of ocelli, the photoreceptive organs used for flight stabilisation in winged insects, which might provide further information for interpreting flight evolution. Our findings support an ancestral loss of wings and that the ancestors of most major lineages were wingless. While the evolution of ocelli was estimated to be dependent on the presence of (fully-developed) wings, ocelli are nevertheless absent in the majority of all examined winged species and only appear in the members of few subordinate clades, albeit winged and volant taxa are found in every lineage. The disjunct distribution of ocelli substantiates the hypothesis on trait reacquisition and that wings were regained in Phasmatodea.

## Introduction

Active flight is considered the key innovation to have driven lineage diversification in animals and has allowed insects to become the most species-rich group on Earth^1–3^. The evolution of the insect wing ∼400 million years ago (mya)^2,4^ and the associated enhanced abilities to disperse and access unreachable resources are directly linked to the remarkable radiation and success of winged insects (Pterygota). Moreover, wings have undergone various modifications to additionally or alternatively serve other functions such as thermoregulation, mate choice and courtship, crypsis and defensive strategies^1,5^. However, despite the numerous advantages, wings have been repeatedly reduced across all pterygote groups with several lineages being completely wingless such as heel-walkers (Mantophasmatodea), lice (Phthiraptera) or fleas (Siphonaptera)^1,6–8^. Paradoxically, also the loss of flight is proposed to be in direct correlation with increased diversification rates and thus also a driver of speciation^9–12^.

Flightlessness appears to occur when the selection for aerial dispersal is relaxed as in habitats with environmental stability or with unfavourable conditions for flight such as high winds or cold temperatures^6,13,14^. As a consequence, flightless taxa may show increased fecundity because resources previously invested in maintaining the energetically costly flight apparatus can now be allocated to reproduction^6,15,16^. The underlying trade-off between dispersal and reproduction has been repeatedly demonstrated for females^17–21^ and males^22–24^, albeit the loss of flight is generally more common in females, often resulting in wing dimorphic species with volant males^1,6,25^. In some species, the dilemma of dispersal capability versus fecundity is solved by wing polyphenism, where an either winged or flightless phenotype is adopted in response to specific environmental triggers^1,16,18,26–29^. Yet, flightlessness is not tantamount to the complete loss of wings (aptery): A phenotype or species may exhibit shortened wing length or retain fully-sized wings but with reduced flight musculature^26,27^.

Within winged insects, the plant-mimicking lineage of stick and leaf insects (Phasmatodea) has been recognised as an expedient study system to investigate the evolution of flight due to their high diversity in wing states and sizes varying among closely related species and between sexes^30,31^ (Figure 1). Generally, forewings are modified to abbreviated and sclerotised tegmina or wing pads, while the hindwings are membranous and folded neatly against the elongated and often slender body at rest^2^. However, most of these herbivorous insects are flightless^1,30^, being either completely wingless (apterous; Figures 1A and 1B) or short-winged (micropterous/brachypterous; Figures 1C–1F), while long-winged forms (macropterous; Figures 1G and 1H) may or may not be capable of ascending flight. Wings that do not sustain powered flight may serve for gliding or other derived utilities such as defensive stridulation or startle displays^32–34^. The latter is common in both short- and long-winged species whose wing undersides or membranes show bright warning colours in contrast to the otherwise inconspicuously coloured body maintaining crypsis (Figures 1E–1H). Sexual wing dimorphism occurs throughout the phasmatodean lineage and is exclusively female-biased^30,31^ (Figure 1I) with the exception of two *Phasmotaenia* species^35^. As a result, females tend to be larger and brachypterous or apterous, whereas males are smaller and either share the female’s wing state or have more developed wings.

**Figure 1.**
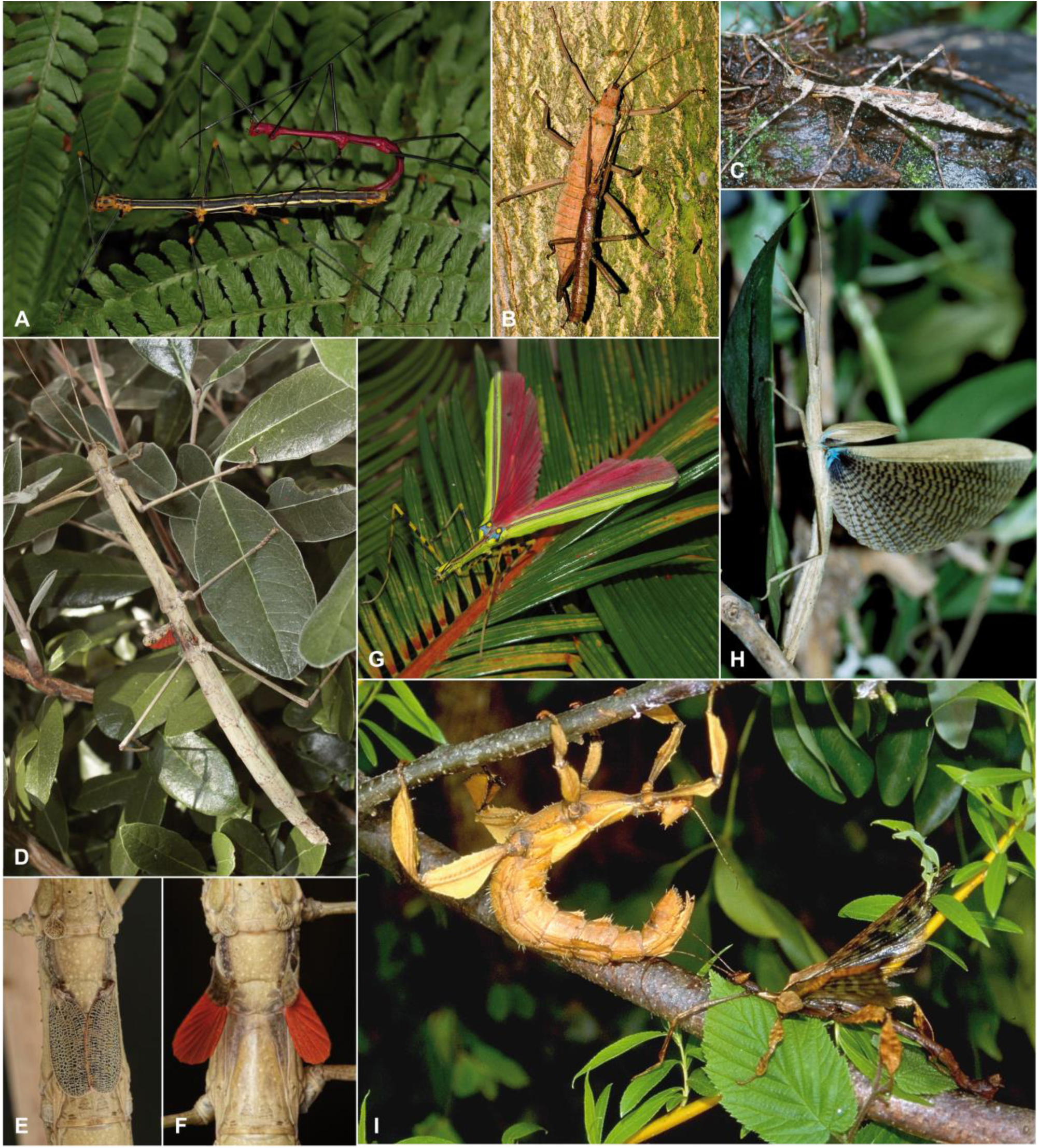
Photographs of various phasmatodean representatives with different wing states: wingless (apterous), short-winged and flightless (micropterous/brachypterous) and long-winged and volant (macropterous). (A) Wingless couple of *Oreophoetes peruana*. (B) Wingless couple of *Eurycantha insularis*. (C) Female of *Pseudodiacantha macklotti* (brachypterous). (D) Female of *Phaenopharos khaoyaiensis* (micropterous/brachypterous). (E and F) Close-up of winglets of *Phaenopharos* sp. (micropterous/brachypterous). The conspicuous colouration is only visible when wings are opened presenting a startle display. (G) Female of *Anarchodes annulipes* (macropterous). The wing membranes exhibit a warning colouration to be used in startle displays. (H) Female of *Metriophasma diocles* (macropterous). The opened wings show the long hindwing and the for phasmatodeans typical shortened forewing. (H) Sexual size and wing dimorphism in a couple of *Extatosoma tiaratum* with brachypterous female on the left and macropterous male on the right. Photos by Bruno Kneubühler and Christoph Seiler.

In contrast to the diverse wing states of extant stick and leaf insects, extinct species of stem group phasmatodeans have fully-developed fore and hind wings^36–38^ suggesting multiple and rather recent shifts from fully-winged to short-winged and wingless forms^31^. Yet, these transition events might not be unidirectional. Whiting et al.^30^ proposed that wings had been completely lost in ancestral stick insects and were independently re-acquired in several descendant phasmatodean lineages. This hypothesis had been extensively debated and criticised as an overstatement in regard to the reliability of the inferred topology and the overall probability of wing regain^39,40^, and was largely considered a violation of Dollo’s law under which the loss of complex traits is irreversible (see Gould^41^, but also Collin and Miglietta^42^ and Goldberg and Igić^43^). The recurrence of wings was however also suggested for water striders and fig wasps^44,45^ as was the re-evolution of various traits such as eyes, digits or teeth in other animals^46–50^. In corroboration with Whiting et al.’s^30^ results, a recent work on phasmatodean wing evolution also concluded a wingless or at least flightless ancestral state for stick and leaf insects^51^. However, both studies’ topologies do not comply with other comprehensive phylogenetic analyses based on large datasets in terms of taxa^52^ and genes^53,54^ and thus may be not reliable. Ultimately, only a robust phylogeny can provide the framework necessary for understanding the evolutionary processes underlying character evolution.

In addition to the flight apparatus itself, the specific nature of sensory systems regulating stabilisation reflexes for maintaining balance during flight such as wind-sensitive hairs^55^ and ocelli^56^ might provide further information for interpreting flight evolution. Pterygote insects generally possess a set of three ocelli (single-lens eyes) besides the faceted compound eyes. These photoreceptive organs process information on light levels more rapidly than compound eyes and thus significantly contribute to horizon detection and orientation in three-dimensional space during flight^56–60^, but have also been suggested to be involved in other functions such as, for instance, the circadian rhythm (see Honkanen et al.^61^, Ribi & Zeil^62^ and references therein). The number and organisation of ocelli and their photoreceptors varies across all insect lineages and may be indicative of the adaptation to different selection pressures^62^. While ocelli are generally lacking in secondarily apterous pterygote insects, they may also be absent in lineages of winged taxa such as webspinners^63^ or beetles^64^. In stick and leaf insects, ocelli appear to be associated with the capability of flight and never occur in wingless taxa^65,66^ (Figure 2).

**Figure 2.**
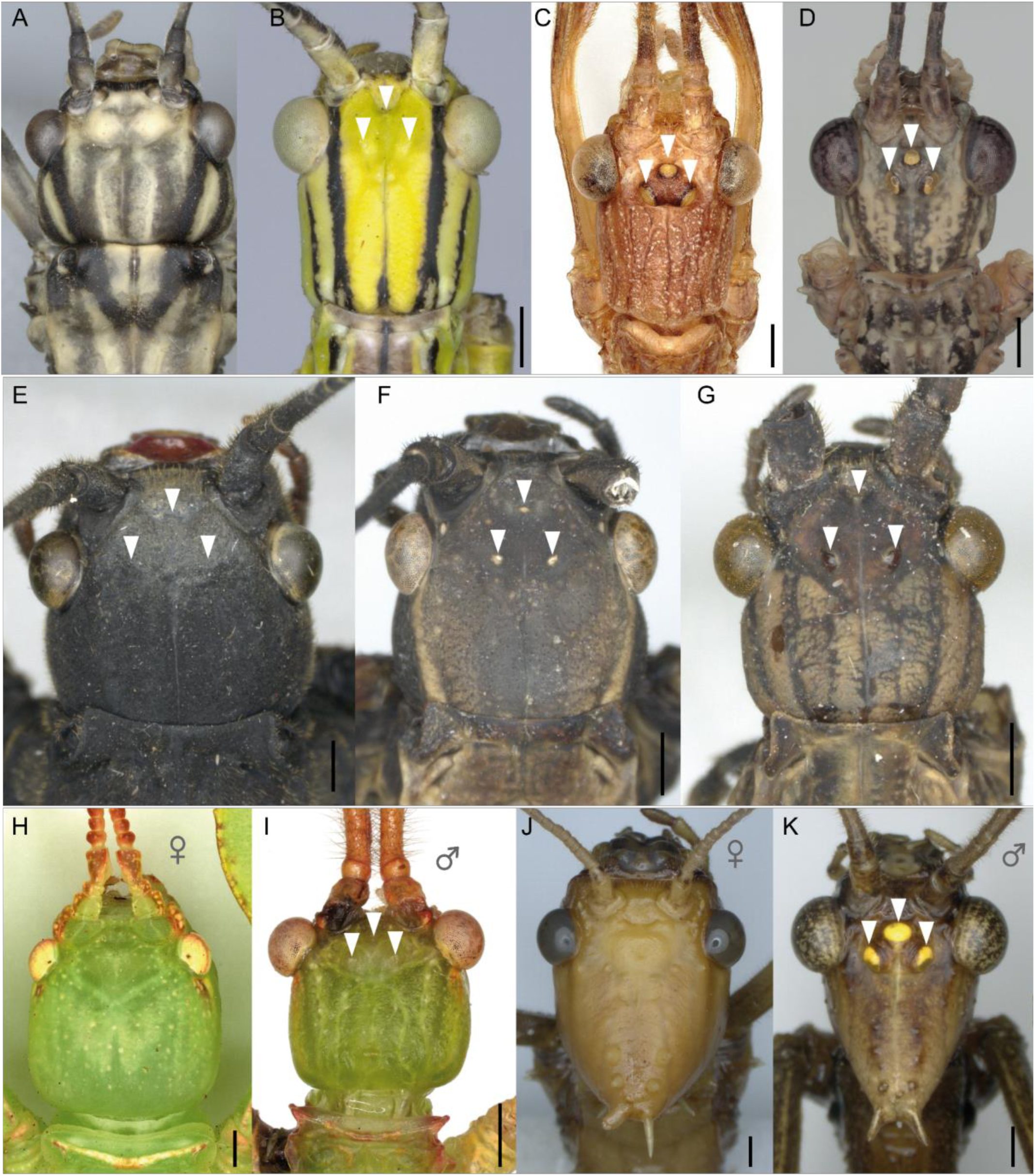
Photographs of the head of taxa with different wing states showing the presence or absence of ocelli. Arrows point to ocelli. (A) Macropterous female of *Aschiphasma annulipes* (Aschiphasmatidae), (B) macropterous female of *Anarchodes annulipes* (Necrosciinae), (C) macropterous male of *Acrophylla titan* (Lanceocercata), (D) macropterous male of *Xeroderus* sp. (Lanceocercata), (E) micropterous female of *Peruphasma schultei* (Pseudophasmatidae), (F) micropterous female of *Pseudophasma scabriusculum* (Pseudophasmatidae), (G) macropterous female of *Pseudophasma fulvum* (Pseudophasmatidae). (H–K) Sexual dimorphism in (H) micropterous female and (I) macropterous male of *Phyllium philippinicum* (Phylliidae), and in (J) micropterous female and (K) macropterous male of *Extatosoma tiaratum* (Lanceocercata). Scale bars: 1 mm. Photos by Tim Lütkemeyer and Marco Niekampf.

Interestingly, numerous winged and volant species lack ocelli nevertheless (Figure 2A). In species that possess ocelli, the degree of their development may reflect whether the species is partially- or fully-winged (Figures 2E–2G). Even within a single species they can be well-developed in the volant male, while completely absent in the flightless female (Figure 2H–2K). Given the assumed strong correlation between flight capability and presence of ocelli, the ocellar system might contribute to differentiate between primarily and secondarily winged stick insects. Herein, we revisit the evolution of flight in stick and leaf insects by inferring a phylogenetic reconstruction of wings and ocelli based on the largest taxon sample to date with >500 species covering all major groups. This extremely dense taxonomic resolution appears to be necessary to satisfactorily reconstruct this highly disparate character system.

## Results

### Phylogenetic signal and ancestral states reconstruction

Morphological data on wings and ocelli were collected for 513 phasmatodean specimens, albeit 18 are only known from one sex (nine females and males, respectively). The majority is found to be apterous with 272 taxa lacking wings entirely and 45 with wingless females (Figure 3). Among winged species, brachyptery is found to be more common in females, whereas males predominantly have fully-developed wings. Ocelli never occur in wingless species and rarely in micropterous taxa (3 of 66 males, 7 of 112 females). Although mostly present in macropterous species, less than half of the examined macropterous females and males possess ocelli. In general, ocelli were found to occur in five of the major lineages, namely, Lanceocercata, Necrosciinae, Pseudophasmatidae, Palophidae and Phylliidae, and females with ocelli exist only in the former three groups.

**Figure 3.**
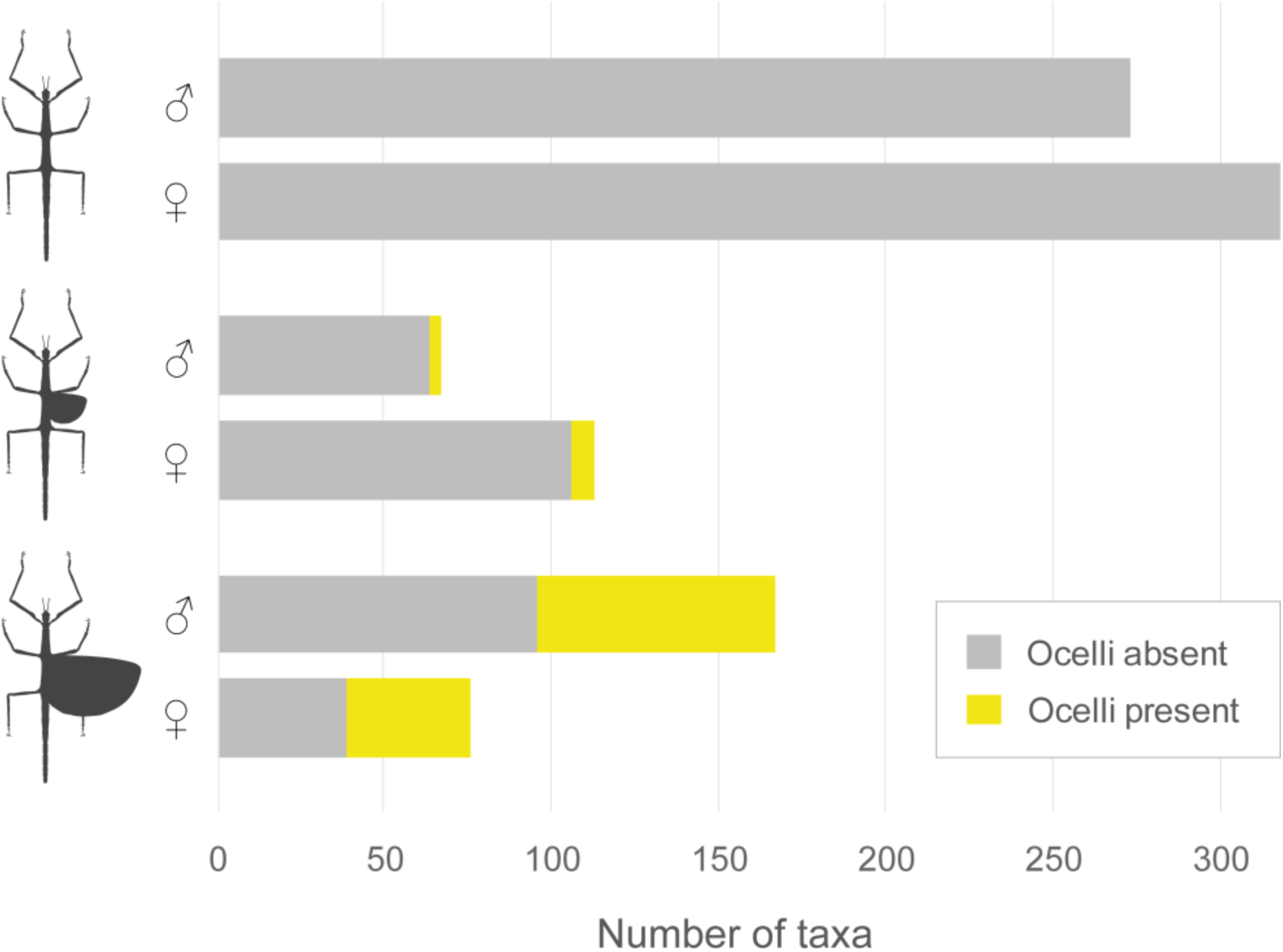
Observed number of wingless, partially-winged (micropterous) and fully-winged (macropterous) male and female phasmatodeans and proportion of taxa with and without ocelli.

A strong phylogenetic signal was detected for all tested traits (presence of ocelli, presence of wings, capability of flight) with negative D statistics (D = -0.568, -0.410 and -0.355, respectively) and fitted lambda values of >0.981 (Supplementary file 6) indicating that these traits are more likely to be shared by closely related species and thus phylogenetically conserved. This was also supported by comparing the numbers of evolutionary transitions observed for each trait (24, 58 and 51) against a randomisation process of that trait (on average 70, 169 and 142; Supplementary file 6).

Prior to reconstructing ancestral states, we fitted different models to the binary wing dataset (data_wings_) and compared log-likelihoods and AIC values (Supplementary file 7). The ARD model was determined as the best-fit model independent of whether the root state was equally probable to be winged/wingless or forced to be wingless. By contrast, the models disallowing the regain of wings (IRR) performed the poorest, even when the root was forced to be winged. Applying the ARD model, the ancestral reconstruction of wings and ocelli estimated the ancestral state for all the major nodes to be wingless and without ocelli for males and females (Figure 4, Figure 4—figure supplement 4 and 5). When regarding the African/Malagasy stick insects as two separate clades, we recognise 21 major lineages of which 14 have winged representatives, but only for 7 the ancestral state was estimated to be winged. The probabilities of ancestral winglessness in males for the higher taxa (Phasmatodea: 48.67%, Euphasmatodea: 84%, Neophasmatodea: 95.33%, Occidophasmata: 96%, Oriophasmata: 97.67%; Figure 4, Supplementary file 8) are not found to be significantly different from the results obtained from the reconstructions based on the phylogenies with different constraints (B1, B3; Figure 4— figure supplement 2 and 3) and in only one instance the macropterous state is estimated to be more likely for the phasmatodean node (B1, Supplementary file 8). The reconstruction of ocelli showed a consistently high likelihood of their absence across the phylogeny and their evolution is estimated to have occurred at least once in Pseudophasmatidae, Phylliidae, Palophidae and Lanceocercata, and three times in Necrosciinae (Figure 4—figure supplement 2–4).

**Figure 4.**
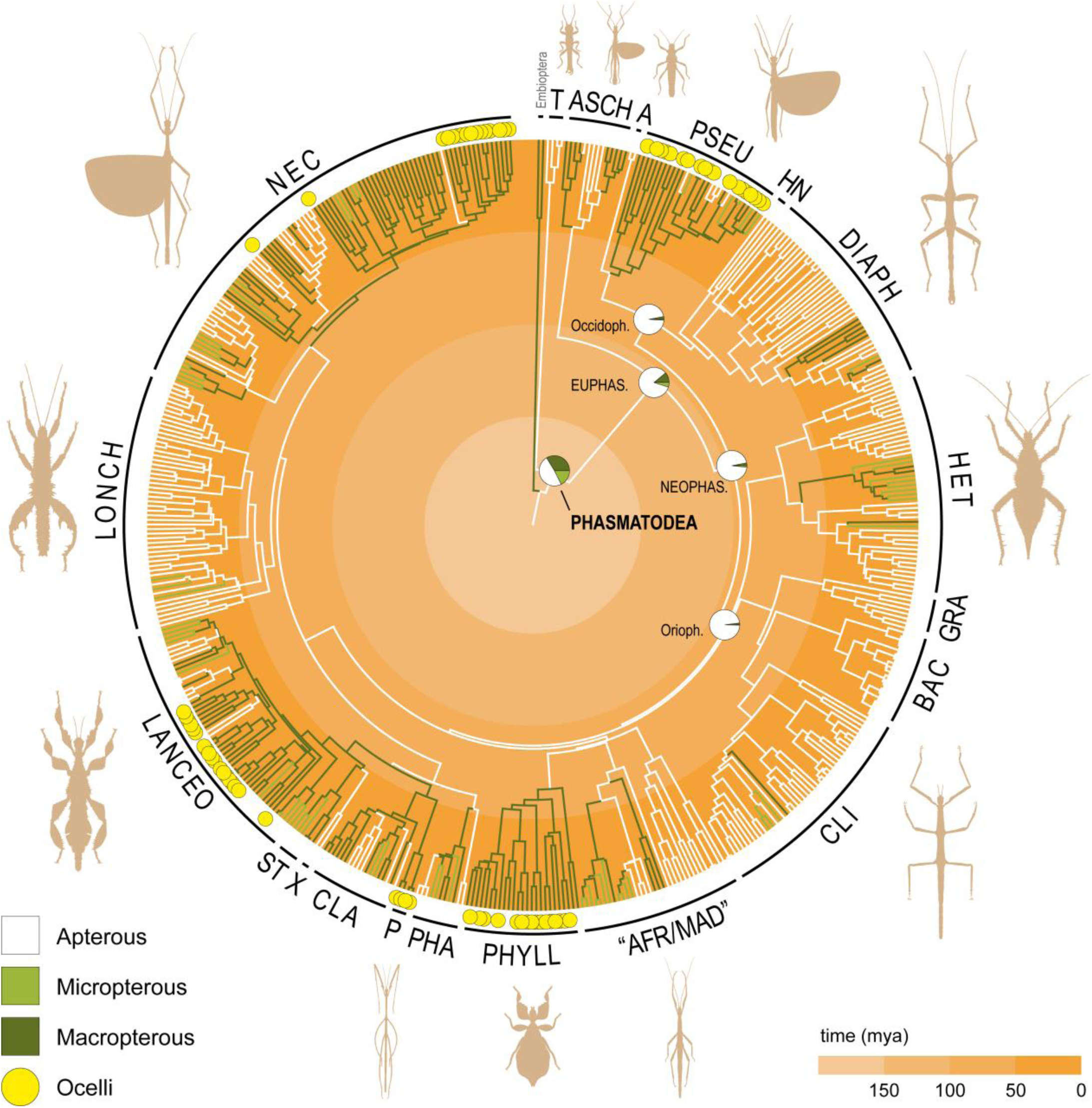
Phasmatodean phylogenetic relationships and reconstruction of wing states. The phylogeny is based on the Bayesian analysis using constraints (B2) and branches are coloured based on the results of the ancestral state reconstruction of male wings (Figure 4—figure supplement 4; see also Figure 4—figure supplement 1–3 and 5). Branch colour for unknown states corresponds to the most likely state of the parent node. Pie charts on major nodes show the probabilities for the ancestral state. The presence of ocelli is highlighted in yellow at the tips. EUPHAS, Euphasmatodea; NEOPHAS, Neophasmatodea; Occidoph, Occidophasmata; Orioph, Oriophasmata; T, Timematodea; ASCH, Aschiphasmatidae; A, Agathemeridae; PSEU, Pseudophasmatidae; HN, Heteronemiinae; DIAPH, Diapheromerinae; HET, Heteropterygidae; GRA, Gratidiidae *sensu* Cliquennois (2020); BAC, Bacillinae *sensu* Cliquennois (2020); CLI, Clitumninae *sensu* Cliquennois (2020); AFR/MAD, African/Malagasy group including Achriopteridae, Anisacanthidae, Antongiliidae, Damasippoididae and Xylicinae *sensu* Cliquennois (2020); PHYLL, Phylliidae; PHA, Pharnaciinae + *Prosentoria*; P, Palophidae; CLA, Cladomorphinae; X, *Xenophasmina*; ST, Stephanacridiini; LANCEO, Lanceocercata; LONCH, Lonchodinae, NEC, Necrosciinae.

The evaluation of the number of transition events between states when wing-regain is permitted (ARD) or wing loss is irreversible (IRR) showed that when analysing the 3-state dataset under the ARD model, the number of shifts was highly increased, albeit most shifts were detected between micropterous and macropterous taxa. Comparing the binary dataset under ARD and IRR models and thus the transitions between the presence and absence of wings revealed more sensible results. Under the ARD model, the loss of wings occurred on average twice as likely as their regain with a total of ∼64 evolutionary shifts on average between states (Figure 5, Supplementary file 9). In contrast, when revolution of wings was not permitted (IRR), the number of losses is significantly higher (∼76; Figure 5, Figure 4—figure supplement 6). Similar results were obtained from comparing the results of ocelli reconstruction with 8 regains and 25 losses under the ARD model compared to 55 losses under the IRR model (Supplementary file 9).

**Figure 5.**
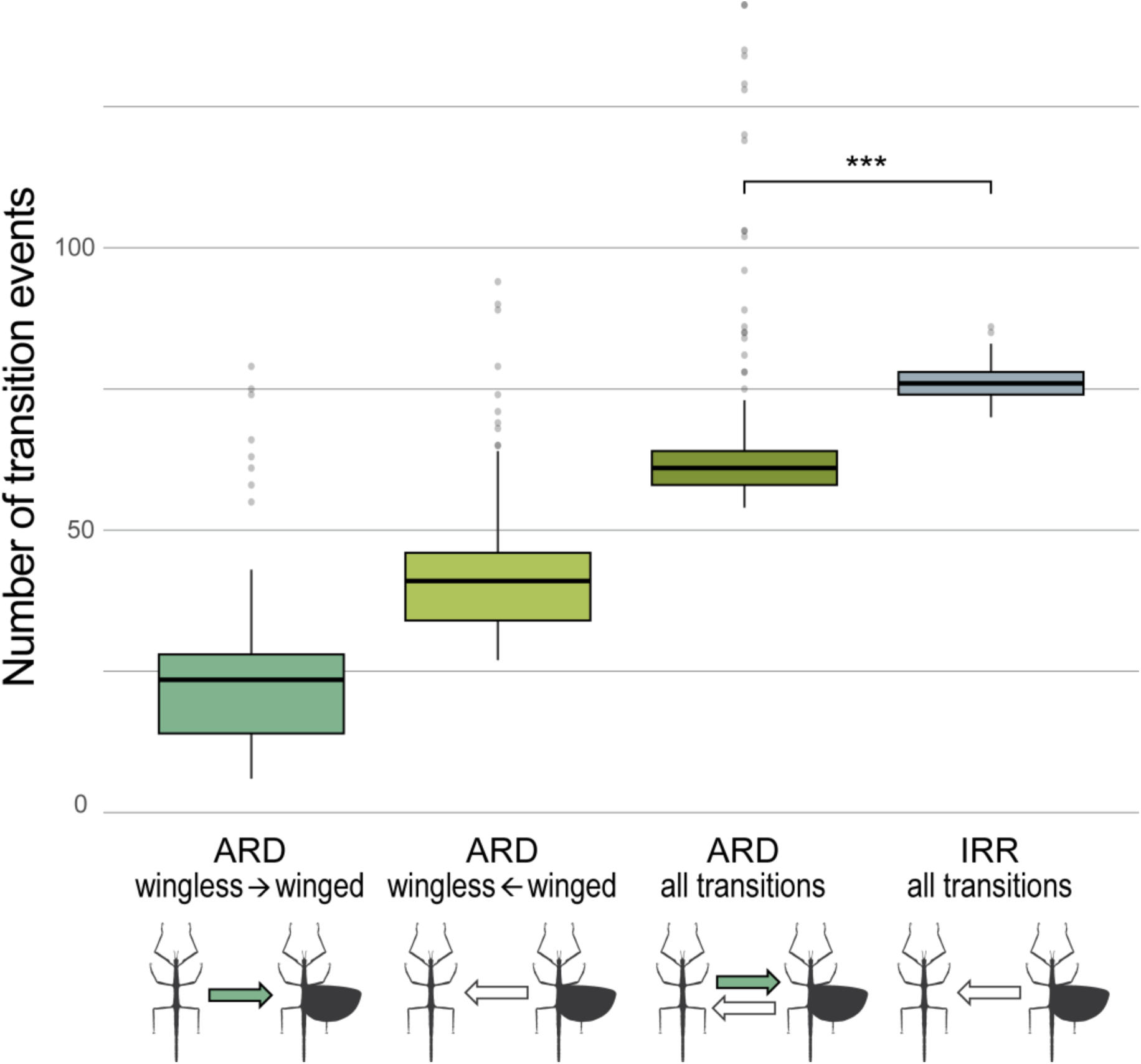
Box-plot diagram of number of transitions between the winged and wingless state. Numbers were generated from performing 300 iterations of stochastic character mapping in phytools based on the binary wing dataset (Figure 5—Source Data 1). The distribution of observed transitions events is displayed as box-plots of the 25th to 75th percentile, with the horizontal line representing the median and vertical lines representing the range (excluding potential outliers). For the ARD model, transition events from wingless to winged and vice versa, and the combined amount are shown. Under the IRR model disallowing wing regain, all transitions are unidirectional. The number of all transitions observed under ARD and IRR is compared using a paired t-test. *** p-value < 2.2e-16.

### Trait correlation

We tested for correlation of wings and ocelli and recovered strong evidence for their correlated evolution in males and females (Supplementary file 10). Specifically, models performed best when ocelli were set as the depending variable implying that the evolution of ocelli was depending on the evolution of wings. However, applying different root states influenced the estimation resulting in varying AIC values and Akaike model weights (AICw). In one instance, when the root state for females was set to wingless + no ocelli, the evolution of ocelli and wings was recovered as intradependent. The wingless root state also resulted in the lowest AICw values for males (AICw = 0.5136), whereas the unconstrained, winged + no ocelli and winged + ocelli root states resulted in higher values (AICw = 0.6462, 0.9434 and 0.7895, respectively). The correlation of wings and ocelli was further examined by fitting models of correlated evolution between ocelli and the three individual wing states. While we did not find any evidence that short wings and ocelli are correlated, there was statistically significant correlation recovered between ocelli and long wings as well as winglessness.

### Diversification rates

The BAMM analysis converged well and ESS values are well above 200. A total of nine diversification rate shifts was detected across the branches of the phasmatodean phylogeny (Supplementary file 11) and the best shift configuration (posterior probability = 0.71) features two shifts that are localised in the clade Euphasmatodea and in the clade of European Bacillinae. The second-best shift configuration additionally shows a slight rate shift for the lineage including among others the Cladomorphinae and Lanceocercata. Overall, the net diversification shows a gradual and constant increase in rate over time and remains comparably low only for Timematodea (Supplementary file 11). The results of our trait-dependent diversification rate analysis to assess the influence of wings on the diversification process show that character-independent HiSSE (CID-4) models were inferred to best fit our data (Supplementary file 12). The best-fit model (AICw = 0.9412), the CID-4 model with four hidden states, does not include the focal character (wings) and thus assumes that changes in diversification rate are independent from the presence or absence of wings.

## Discussion

Independent of the alternative backbone phylogenies (for more details see Appendix 1), the reconstruction of the two key characters involved in insect flight produced almost identical results in all analyses (Figure 4, Figure 4—figure supplement 2–4, Appendix—figure 1). Applying the ARD model and thus allowing transitions rates to be different for gains and losses, the ancestral reconstruction of wings and ocelli estimated the ancestral state to be wingless and anocellate for the major nodes of male and female phasmatodeans (Figure 4, Figure 4—figure supplement 4 and 5). Although *Timema* and their extinct relatives are completely wingless, the node at the split between Timematodea and Euphasmatodea is however not estimated to be unequivocally in favour of a wingless ancestral form. Under consideration of the significant time gap of approximately more than 50 million years between their divergence and the start of euphasmatodean diversification that has repeatedly been inferred^4,51–54,67,68^, it cannot be ruled out that stem group euphasmatodeans were winged. The scarce fossil record^66,69,70^ shows that extinct Phasmatodea from the Cretaceous were predominantly winged^36–38,71–73^, but are thought to belong to the stem group of Phasmatodea rather than Euphasmatodea^66,70,73^. Hence, the fossil evidence is not contradicting a wingless ancestor of crown Phasmatodea or Euphasmatodea. Moreover, it is noteworthy that the wing morphology of these Mesozoic winged stick insects is different from that of recent phasmatodeans^38,74^: All fossil specimens exhibit two long pairs of wings^36–38^, whereas the sclerotized forewings (tegmina) of extant macropterous forms are shortened. Long tegmina are in fact considered the plesiomorphic condition of Polyneoptera, which are suggested to have been independently reduced in various lineages such as in stoneflies or webspinners as well as stick and leaf insects^75^, albeit it remains unclear when and how often forewing shortening occurred in Phasmatodea. Also the fossils’ tegmina venation is different and its modification in modern phasmatodeans might be due to a convergent evolutionary process of wing shortening and not because of common ancestry, especially when considering the few extant, unrelated and strongly subordinated species that exhibit long tegmina with venation differing from other recent forms (i.e., *Heteropteryx, Prisopus*, Phylliidae)^74,76,77^. While the large tegmina of the flightless leaf insect females (Phylliidae) contribute to their leaf mimicry and therefore might have been secondarily elongated^78,79^ (the volant males possess shorter tegmina), the wing morphology of male *Heteropteryx* is most similar to that of stem phasmatodeans^36,74,77^. As subordinated lineage within the ground-dwelling and predominantly wingless Heteropterygidae, it was suggested that *Heteropteryx* secondarily became arboreal and the male regained the capability of active flight. Due to the most plesiomorphic wing morphology among extant stick and leaf insects, its wings were hypothesised to be the product of an atavistic reversion^66,76^. Our results support this hypothesis and not only for this clade, where morphology is corroborative, but also for the more diverse winged lineages of Euphasmatodea.

Regardless of whether wings were present or absent in the common ancestor of Euphasmatodea, our results show an increase in diversification rate with the start of their radiation (Supplementary file 11). This rapid radiation that largely follows the Cretaceous-Palaeogene (K-Pg) mass extinction (∼66 mya) was previously suggested to be linked to the diversification of flowering plants^53^ – a co-evolutionary pattern also observed in other plant-associated insect groups^80–82^. Our results substantiate this hypothesis, particularly, when considering the results of the trait-dependent diversification analysis (HiSSE) from which we concluded that flight or flightlessness might not have been the main driver of euphasmatodean diversification. Alternatively, the evolution of hard-shelled eggs^52,77,83^ and the acquisition of endogenous pectinase genes^84^ further shaping the co-evolution with their food plants are most likely to have contributed to the success of the early euphasmatodean lineages. Also the recurrent opportunities for colonising new land masses (i.a., the Indo-Pacific region) appears to have promoted speciation^76,85^ and might explain the increased diversification rate recovered for Lonchodidae, Lanceocercata and relatives within Oriophasmata (square symbol, Supplementary file 11). By contrast, the slight decrease in the rate for the European Bacillinae (circle symbol) might be an artefact resulting from oversampling closely related taxa, most of which are only represented by one single gene, but could also correspond to the ecological shift to a temperate region. Given that both the presence of wings and the lack thereof were shown to potentially increase diversification in insects^1,2,6,9–12^, it cannot be excluded that either condition had an equal impact on the diversification of the individual euphasmatodean lineages (e.g., for the predominantly macropterous and most species-rich lineage Necrosciinae versus the speciose and mostly wingless Lonchodinae^66,86,87^). Ultimately, that no significant change in rate among subgroups was detected and character-independent diversification was favoured is indicative of the wing state not being subject to Dollo’s law of irreversible evolution^43^.

Within Euphasmatodea, only few lineages were recovered as ancestrally winged (e.g., Pseudophasmatidae, Phylliidae), whereas in others, the winged taxa are found rather subordinated among otherwise wingless forms (e.g., Lonchodinae, Diapheromerinae). In contrast to the seemingly random distribution of apterous species in ancestrally winged lineages, the winged species in predominantly wingless groups are closely related, indicating single origins of wing re-evolution. Discrepancies may be explained by incomplete taxon sampling or wrongly recovered phylogenetic position, which might diminish in a phylogenomic context or when applying an even denser taxon sampling. Conversely, the multiple instances of secondarily apterous taxa found among winged lineages are the result of the higher probability of wing loss in comparison to that of regain^30,39,40,45^. Generally, evolutionary transitions were proposed to occur rapidly and often in Phasmatodea – not only between apterous and macropterous taxa but also between wing states^31^. Previous authors argued that the probability or cost ratio of wing gain was considered too high compared with that of its loss^40^. In our reconstruction, the estimated number of potential regains ranges between 9 and 36 events (∼22) with ∼42 losses (27–59) versus ∼76 losses (71–82) if wing loss is considered irreversible (Figure 5), hence, permitting re-evolution appears to be more parsimonious. For instance, in predominantly wingless Diapheromerinae, we find 1–3 potential wing regain events versus an alternative of a total of ten losses if wing recovery was impossible. Although our results clearly support the hypothesis of re-evolution, we nevertheless acknowledge that the reacquisition of a complex trait such as wings must be assumed less likely than its loss, and that it is possible that ultimately, independent wing loss might have occurred much more often in phasmatodeans than in other insect groups. However, how can we then explain the distant disjunct distribution of ocelli in Phasmatodea?

Insect ocelli have not been intensively studied in a phylogenetic context before, but it is well-known that there are winged lineages within Pterygota that lack ocelli such as earwigs (Dermaptera) and webspinners (Embioptera). Although both groups have undergone a unique modification of wings, their specific lifestyle led to a reduced need for aerial dispersal over time (ground-dwelling lifestyle of Dermaptera; silk galleries of Embioptera)^75,88,89^, which generally resulted in 20–40% of flightlessness and the complete loss of wings in all embiopteran females^1^ and potentially promoting the loss of ocelli. In contrast, the majority of phasmatodean taxa are flightless or wingless^1,30,66^, and some winged species do possess ocelli (Figure 3). Considering the significant number of macropterous anocellate species and the seemingly arbitrary distribution of ocelli-bearing lineages raises the question whether ocelli might have re-evolved; otherwise, if ocelli had been in the ground plan of winged (Eu)Phasmatodea, why would most lineages that retained the capability of flight have reduced this flight-stabilising sensory system? For instance, the Aschiphasmatidae, sister group to the remaining Euphasmatodea (=Neophasmatodea), comprise besides wingless forms several fully-winged species that are capable of active flight, yet, none possess ocelli. Our results show that the presence of ocelli is restricted to five subordinated and completely unrelated lineages, albeit winged and volant taxa are found more frequently across Phasmatodea (Figure 4). While it is plausible to assume that ocelli might have been lost in winged lineages that contemporarily lost the capability of flight (i.e., micropterous lineages), the high number of macropterous and volant species lacking ocelli is perplexing. The alternative scenario involving the possibility of wing regain implies that secondarily winged lineages re-evolved the ocelli subsequent to or simultaneously with the recovery wings, which is highly supported by our analyses. Specifically, the ancestral state of Pseudophasmatidae, Palophidae and Phylliidae was recovered as possessing ocelli along with wings, whereas ocelli are estimated to have occurred only in subordinated clades in Lanceocercata and Necrosciinae in spite of wings and flight being recovered to have a more ancestral. The necrosciine taxa represent an optimal example, since ocelli occur only in one single subclade within a highly diverse macropterous lineage and are otherwise only present in two unrelated species, where they must have re-appeared independently (*Hemisosibia incerta, Gargantuoidea triumphalis*). Conversely, it might be a common trend for brachypterous species within the ocelli-bearing clades to reduce ocelli (as observed in other insect groups^90–93^), albeit there are also few macropterous species lacking ocelli (e.g., *Creoxylus, Eurynecroscia, Paraprisopus*).

In comparison to the other ocelli-bearing clades, the distribution of ocelli within Phylliidae appears more ambiguous. While females are sedentary and flightless, all males are volant and depend on flight for mate localisation^66,85,94^, thus, it is questionable whether there have been multiple independent secondary losses of ocelli as estimated by our analysis. Particularly the phylogenetically incoherent degree of ocelli development in the species of *Phyllium*, where ocelli may be absent or weakly, moderately or well developed (not coded in our analysis), suggests that the ocellar system is a disparate character, which might have gradually and independently re-evolved in several phylliid lineages. The possibility of ocelli reacquisition is further supported by photographs^95^ and specimens that we examined of the Giant Malaysian Leaf Insect (*Pulchriphyllium giganteum*) showing the presence of ocelli not only in males, but also in females. This detail is lacking from the original species description^96^ and from subsequent morphological studies including this species^97,98^. Contradicting the previous assumption that female leaf insects are entirely devoid of ocelli^66,99^, this discovery may be interpreted as further evidence for secondary ocelli gain – even in flightless species or sexes. Generally however, it cannot be excluded that a potentially functional ocellar system may be present internally in the absence of an external (visible) lens as it has been observed in other insects^100–104^ and that this condition preceded the development of fully-formed ocelli at least in some lineages such as *Phyllium*. The alternative scenario involving (gradual) reduction of ocelli in volant stick and leaf insects does not appear plausible especially under consideration of the different functions of ocelli besides those related to flight^57,58,61,62,105^. Also the predominantly nocturnal life style would rather promote the enlargement of ocelli and not their reduction^58,106^. Similarly, it is also argued that in dense vegetation, where there is no clearly visible horizon line, flying insects such as Necrosciinae would undergo specific modifications to improve the performance of the ocellar system to compensate for the obscured horizon^62,105^, yet, there are no habitat differences between ocellate and anocellate Necrosciinae. Hence, we favour the hypothesis that ocelli are not plesiomorphic with copious losses but instead were re-evolved as was also proposed for lineages of the predominantly anocellate beetles^64,107^. To further corroborate this assumption, future studies are needed to elucidate the organisation and development of ocelli across phasmatodean taxa as well as to examine the internal morphology of the head capsule to clarify the potential presence of internal structures in anocellate species that facilitate ocelli emergence.

Under the assumption of Dollo’s law of irreversible evolution, the developmental genetic foundation of a complex trait is lost along with the trait itself and therefore cannot be reacquired because evolving the same complex structures *de novo* is not considered possible. However, if the underlying genetic architecture for the lost trait remains conserved, reinitiation of suppressed genes might result in its recovery. Moreover, evolutionary changes in gene regulatory networks are proposed to have a major influence on morphological evolution^108,109^. Given that the genetic components and developmental processes of a complex character were conserved, evolutionary shifts in developmental gene expression patterns might be responsible for the loss as well as the regain of a trait^108^. The key elements underlying insect flight have been conserved over hundreds of million years of evolution and may remain available for reactivation in secondarily wingless insects for a long period of time^110^, albeit it was estimated that silenced genetic pathways cannot retain function for more than 10 million years^111^. Yet, regulatory elements may be reinitiatable over longer periods of time^50^, especially when the genetic developmental programme is underlying pleiotropic maintenance and is accommodated in similar structures or tissues still present^30,47,48,50,108^. Since the genetic basis for insect wing development is retained in leg formation^112,113^, genes may have been co-opted from these existing expression patterns that then elicited wing re-evolution in stick and leaf insects^30^. Although a gradual process of the reversal of wing loss involving the necessary deployment of multiple genes must be assumed, it has been shown that musculature and innervation associated with flight can be maintained in flightless phasmatodeans^114^, while in other insects flight loss may be explained by degradation of these structures^27^. Therefore, we suggest that in phasmatodeans apparently a latent and hitherto not understood capacity to produce wings and flight-associated traits exists (or existed) that facilitated the reappearance of wings, flight and ocelli to occur multiple times independently. Our hypothesis thus incorporates that the disjunct distribution of these traits did not result from plesiomorphy and numerous losses but from a radiation of wingless ancestors whose descendants independently regained wings (and ocelli). This is further supported by the effects attributed to the loss of flight in the wingless ancestor such as increase in body size and reduction in eye size^1,57,115^ of which the latter is still observable in modern volant species indicating a rather recent regain of flight. Since the advantages of both flight and flightlessness are equally abundant, the trade-off between dispersal and fecundity is reflected in phasmatodean capability of the dynamic transition between wing states as well as by the high occurrence of sexual dimorphism in terms of size and wings^31^.

The currently expanding body of research reporting re-evolution and reversals clearly challenges our assumptions about the irreversibility of complex traits. Our study contributes to this development and establishes a basis for future work to further investigate the regain of wings and flight in stick and leaf insects including the comparison of wing lengths as a continuous character and the assessment of flight capability. Ideally, morphological, developmental, phylogenomic and palaeontological work can eventually be united into a comprehensive picture to understand phasmatodean wing evolution and the underlying genetic developmental processes that lead to the manifestation of atavistic reversals.

## Material and methods

### Taxon and gene sampling

Our comprehensive dataset consisted of 513 species and specimens with representatives of all major clades of Phasmatodea and covering almost 50% of its generic diversity (Supplementary file 13). For the outgroup, we further included two species of Embioptera, which were repeatedly recovered as the sister group of Phasmatodea^4,30,53,116^. Our gene sampling comprised three nuclear (18S, 28S, H3) and four mitochondrial loci (12S, 16S, COI, COII) of which data for numerous specimens were already available on GenBank^30,51,52,67,76,85,86,97,117–126^. Additionally, we generated new data for 194 specimens following the protocol given by Bank et al.^76^ with primers adopted from Buckley et al.^117^ and Robertson et al.^52^ (see Bank et al.^76^) and deposited the sequences in GenBank (Supplementary file 13).

### Phylogenetic analysis and divergence time estimation

Phylogenetic comparative analyses are highly dependent on a robust phylogeny. Hence, in order to optimise our topology, we modified our tree inferences to mimic the phylogenetic relationships obtained from transcriptomic studies^53,54^ by constraining the higher taxonomical groups (i.e., Phasmatodea, Euphasmatodea, Neophasmatodea, Oriophasmata and Occidophasmata).

The final supermatrix of 5636 bp (Supplementary file 14) was obtained by aligning, trimming and concatenating the sequences as described by Bank et al.^76^. We partitioned the dataset by gene and by codon position for ribosomal (12S, 16S, 18S, 28S) and protein-coding (COI, COII, H3) genes, respectively, prior to determining the best-fit partitioning scheme and evolutionary model using ModelFinder^127^ (options -m MF --merge greedy) integrated in IQ-TREE v. 2.1.2^128^. The third codon position of the COI and COII genes as well as the second and third position of H3 was merged, resulting in 11 partitions (see Supplementary file 15).

We applied two approaches to infer the phylogenetic relationships: Maximum Likelihood (ML) and Bayesian Inference (BI). For the ML tree inference, we used IQ-TREE^128,129^ to conduct 100 independent tree searches using the aforementioned constraints and the best-fit partitioning scheme and substitution models. For each search, we used a random starting tree, increased the number of unsuccessful iterations (-nstop 500) and set the perturbation strength to 0.2. We computed the log-likelihoods and at the same time tested the tree topologies to identify the best-scoring tree. Nodal support was subsequently calculated using the Ultrafast Bootstrap approximation (UFBoot) and the SH approximate likelihood ratio test (aLRT)^130^ by generating 10,000 replicates. UFBoot support values were plotted on the best tree, whereas for the SH-aLRT test, a new phylogeny was generated by default in IQ-TREE.

For the BI tree, we simultaneously conducted the phylogenetic analysis with the divergence time estimation in BEAST2 v. 2.6.1^131^. We used the same partitioning scheme as for the ML analysis, but substitution models were selected by bModelTest v. 1.2.1^132^ implemented in BEAST. We linked trees and clocks across all partitions and applied the Yule tree prior and a relaxed clock with an uncorrelated lognormal distribution (UCLD) and a clock-rate of 1e^-7^. We time-calibrated the tree using five fossils unambiguously assigned to Phasmatodea (Supplementary file 16) and chose a lognormal prior distribution with the minimum age selected as the offset and log-mean and log-sd set to 1.0. In the case of the two fossils from Dominican amber (*Clonistria* sp. and *Malacomorpha* sp.), the calibrated node was selected to include the sister group (option “use originate”). For these fossils as well as for the Phasmatodea calibration, the log-mean was set to 2.0 to allow a softer maximum bound. To account for potential phylogenetic incongruences that might have an impact on the reconstruction of ancestral states, we performed three independent analyses based on different constraints: For the first, we used the same constraints as for the ML analysis (B1), for the second we additionally constrained the Heteropterygidae as the sister group to all the remaining Oriophasmata (following the results of transcriptomic studies^53,54^) (B2), and for the third, we did not set any constraints for Oriophasmata and Occidophasmata and thus only for Phasmatodea, Euphasmatodea and Neophasmatodea (B3). All analyses were run for 300 million generations with parameter and tree sampling every 10,000 generations. Convergence was assessed in Tracer v. 1.7.1^133^ and a Maximum Clade Credibility tree was generated after removing the first 3000 trees as burn-in.

### Phylogenetic comparative methods/Comparative analysis/Diversification analysis

To compile our morphological data matrix, we gathered information on the wing states and the possession of ocelli for both sexes for each taxon included (Supplementary file 17). For specimens not available for examination, we reviewed the literature (e.g., Redtenbacher, 1906, 1908) and searched for suitable photographs of wild, captive-bred or type species online (www.phasmida.speciesfile.org, www.phasmatodea.com). While we assembled the ocelli data based on their presence or absence and did not distinguish between different types of ocelli (e.g., number, relative size), wing states were coded as wingless (apterous, 0), partially-winged (micropterous, 1) or fully-winged (macropterous, 2). The differentiation between micropterous and macropterous was made arbitrarily as we defined a morph as fully-winged when the hind wings exceeded the fourth abdominal segment^77^. Since some analyses require binary data, we created two additional datasets by combining the three wing states based on wingless (0) versus winged (1) taxa (data_wings_) and based on flightless (0) versus fully-winged (1) taxa (data_flight_). Missing data was generally coded as absent (0). For all analyses, the fossil-calibrated BEAST tree with B2 constraints was used.

### Phylogenetic signal

Using the binary datasets for males, we estimated the phylogenetic signal of the presence of ocelli, presence of wings (data_wings_) and capability of flight (data_flight_) by calculating the D statistic^136^ in the R package caper v.1.0.1^137^ using the phylo.d function with default parameters and 10,000 permutations. Furthermore, we assessed phylogenetic signal using the function fitDiscrete in the R package geiger v.2.0.7^138^ to fit Pagel’s lambda^139^ under the ARD transition model. We expected D estimates to be ≤0 and lambda values closer to 1, both serving as an indicator for traits having evolved under a Brownian motion model and not to be recovered as randomly distributed across the phylogeny. To further evaluate a potential random distribution, we created a randomised character matrix to compare the number of evolutionary transitions with the results from our true dataset as outlined elsewhere^140,141^.

### Ancestral state reconstruction

Ancestral states of wings and ocelli were estimated in the R package phytools v.0.7-70^142^. To determine the best-fit model, we used the functions fitMk and fitDiscrete to fit different models to the binary dataset of male wing traits (datawings). Using fitDiscrete, we tested models that permitted evolutionary rates between wingless and winged to be equal in both directions (equal-rates, ER), different in all directions (all-rates-different, ARD) and unidirectional from winged to wingless (customised irreversible model, IRR). We repeated the tests using the function fitMk and additionally tested the IRR model with the root state fixed to “winged” and the ARD model with the root state fixed to “wingless”. Finally, for the ancestral state reconstruction, we used the 3-states datasets of wings and the binary datasets of ocelli for both males and females, and coded missing data with equal probabilities for each of the possible states. We performed stochastic character mapping^143^ with the function make.simmap implemented in phytools based on the best-fit model (ARD) and ran 300 iterations of Markov chain Monte Carlo (MCMC) sampling. Using the option Q=“mcmc”, the rate matrix Q is sampled 300 times from its posterior probability distribution using MCMC and stochastic maps and node states are generated conditioned on each sampled value of Q. To assess whether the topology of the tree had an impact on the estimated ancestral states, we also performed the analysis using the alternative trees (B1 and B3 constraints).

Given the 300 inferred possible evolutionary histories resulting from stochastic character mapping, we calculated the average number of transition events between wings states and the presence/absence of ocelli. For comparison to an evolutionary hypothesis where wing/ocelli regain is not permitted, we repeated simulated stochastic maps as described above but using the IRR model. To be even more conservative, we also reconstructed ancestral states using both models for the binary wing dataset (data_wings_).

### Trait correlation

We used Pagel’s binary character correlation test^144^ as implemented in phytools to assess potential coevolutionary dynamics between wings and ocelli in males and females. By applying the ARD substitution model and using different parameters for the argument “dep.var”, we estimated whether wings and ocelli evolved independently, intradependently or whether the evolution of wings was dependent on the evolution of ocelli or vice versa, and determined the best model to explain the correlated evolution with the Akaike information criterion (AIC) and Akaike weights. As tests for correlation of traits may be sensitive to the root state, we explored the impact of applying different root states by fixing the root to represent either wingless + no ocelli, winged + no ocelli, and wings + ocelli. We further tested for the correlation of ocelli and either of the three wing states (apterous, micropterous, macropterous). We estimated and compared the values of log-likelihoods and Akaike weights, and corrected p-values for multiple comparisons using the Bonferroni correction.

### Diversification analysis

We explored evolutionary rate dynamics with the Bayesian analysis of macroevolutionary mixtures (BAMM) and the R package BAMMtools^145,146^. Priors were estimated using the BAMMtools function “setBAMMpriors” and modified accordingly (expectedNumberOfShifts = 1.0; lambdaInitPrior = 4.80107993368211; lambdaShiftPrior = 0.00644760145800591; muInitPrior = 4.80107993368211). We ran four chains of 10 million generations being sampled every 1000 generations using the “speciationextinction” model and seting “globalSamplingFraction” to 0.15 (∼500 taxa of ∼3400 described species). Convergence was assessed and subsequent data analysis performed in BAMMtools.

Whether the evolution of wings had an impact on diversification rates was tested using the Hidden State-dependent Speciation and Extinction model (HiSSE) in the R package hisse^147^. We used the binary dataset of male wings and removed the outgroup (Embioptera). Following Song et al. ^148^, we then fitted 24 different models ^147^ including original BiSSE models ^149^, character-independent diversification models (CID-2 and CID-4) and HiSSE models, which assume a hidden state. All models were compared using log-likelihood, AIC and Akaike weights.

## Data and code availability

Newly obtained sequences have been deposited in GenBank under accession numbers xxx–xxx (see also Supplementary file 13). Figure 5—Source Data 1 contains the numerical data used to generate Figure 5. Additional data and code for this paper are available at GitHub (xxx).

## Acknowledgements

We are indebted to the countless people who provided numerous specimens over the last years, with special thanks going to Thomas R. Buckley (New Zealand), Nicolas Cliquennois (Madagascar), Sylvain Hugel (France), Gavin Svenson (USA), Michael F. Whiting (USA), Albert Kang (Malaysia), Bruno Kneubühler (Switzerland), Kristien Rabaey (Belgium), Valerio Scali (Italy), Katja Kramp (Germany), Savel R. Daniels (South Africa) and Donald Windsor (Panama). For information concerning the ocelli presence in certain taxa, we thank Royce T. Cumming (USA), Albert Kang (Malaysia), Nicolas Cliquennois (Madagascar) and Francis Seow-Choen (Singapore). We thank Katharina Henze for helping in the molecular lab. Photos were kindly provided by Bruno Kneubühler, Tim Lütkemeyer, Marco Niekampf and Christoph Seiler. This study was supported by the German Research Foundation (DFG grants BR 2930/4-2, BR 2930/5-1 and BR 2930/7-1).

## Author contributions

Conceptualisation, Bank and Bradler; Funding acquisition, Bradler; Formal analysis, Bank; Investigation, Bank; Visualisation, Bank; Writing – original draft, Bank; Writing – review & editing, Bank and Bradler.

## Figure legends

**Figure supplement 1.**
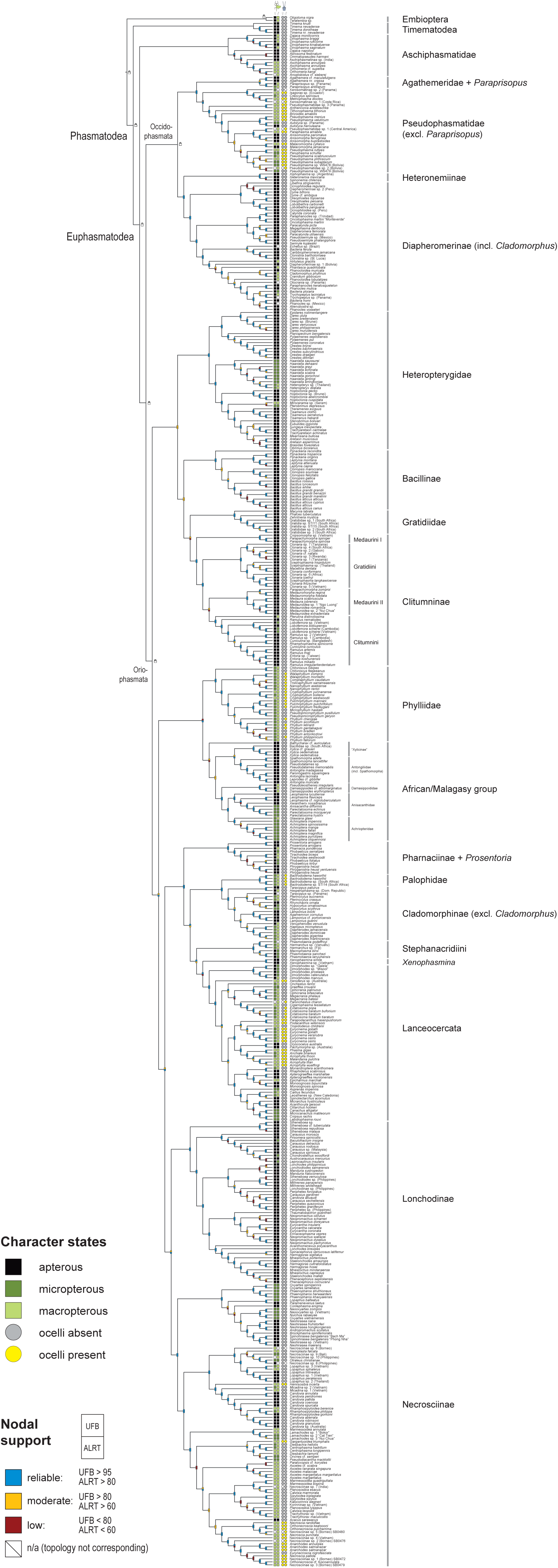
Maximum Likelihood phylogeny based on the best-scoring tree with nodal support (UFBoot and SH-aLRT) at each node (Supplementary files 1 and 2). Lock symbols represent constrained clades (B1 constraints). Character states for wings and ocelli are depicted at tips for females and males.

**Figure supplement 2.**
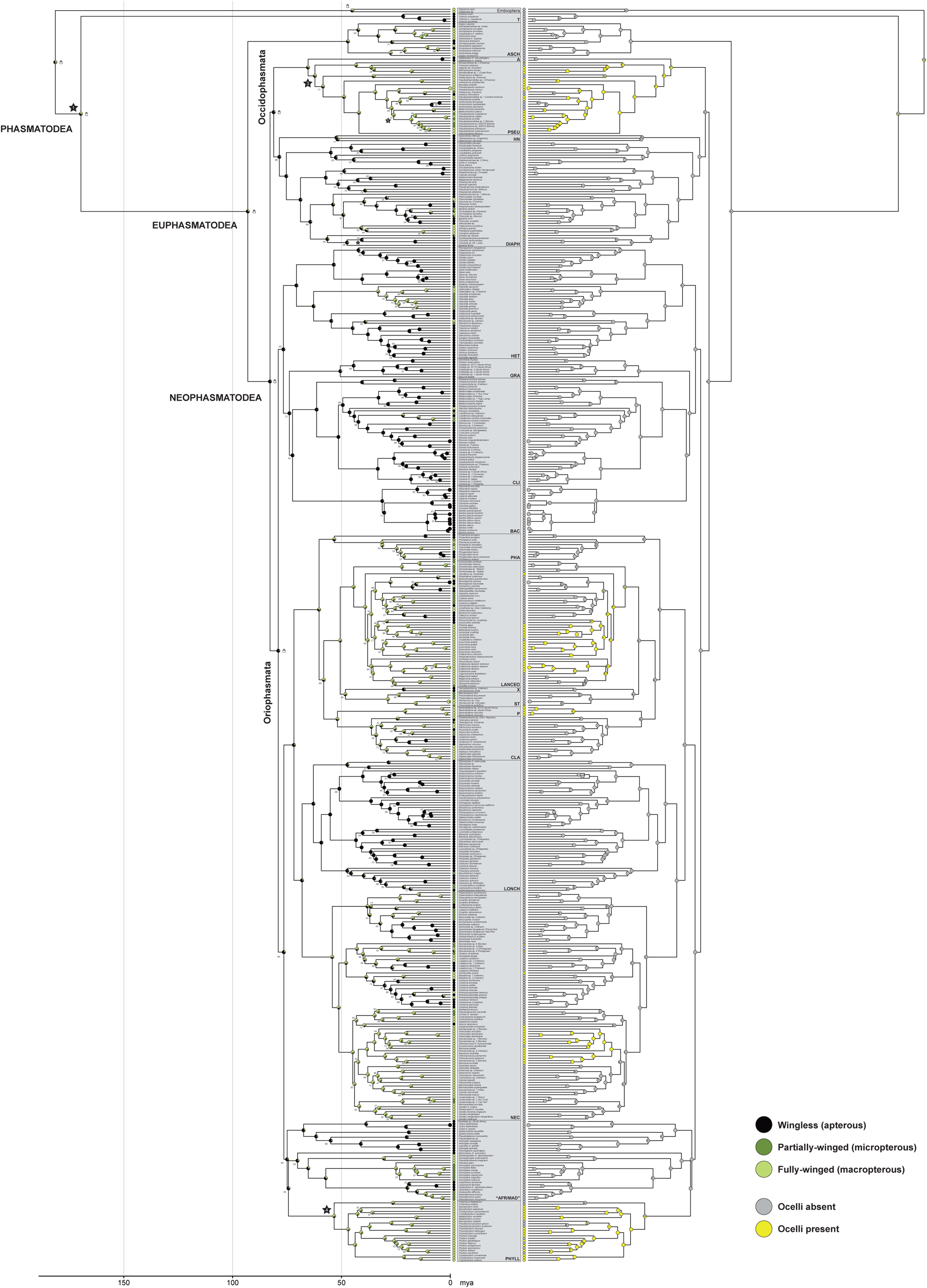
Ancestral state reconstruction for males based on the BI tree with B1 constraints (see lock symbols at nodes; Supplementary file 3). Nodal support values (<1 posterior probability) depicted at each node. Stars represent the fossils used for calibration and numbering corresponds to ; Supplementary file 16.

**Figure supplement 3.**
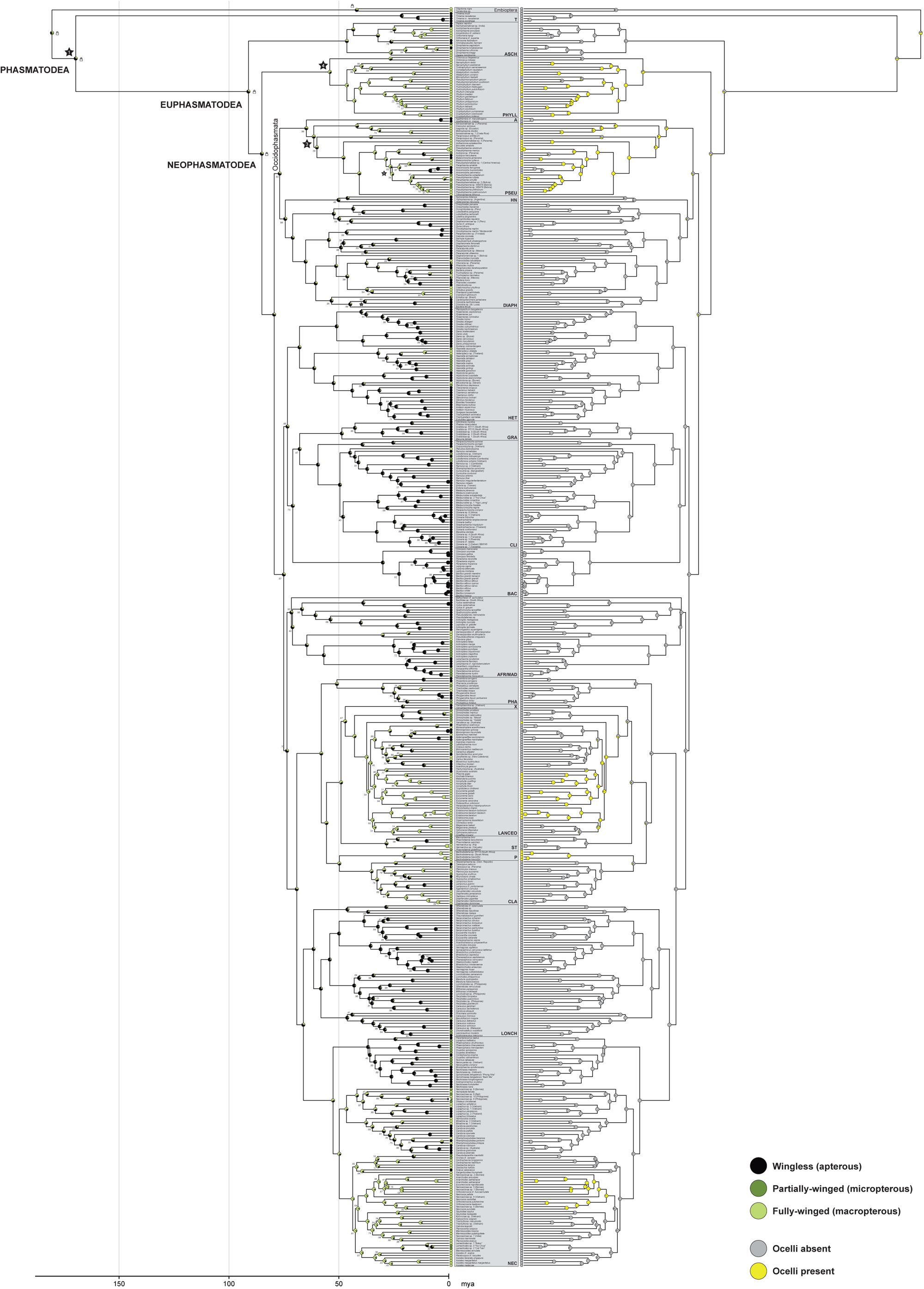
Ancestral state reconstruction for males based on the BI tree with B3 constraints (see lock symbols at nodes; Supplementary file 5). Nodal support values (<1 posterior probability) depicted at each node. Stars represent the fossils used for calibration and numbering corresponds to Supplementary file 16.

**Figure supplement 4.**
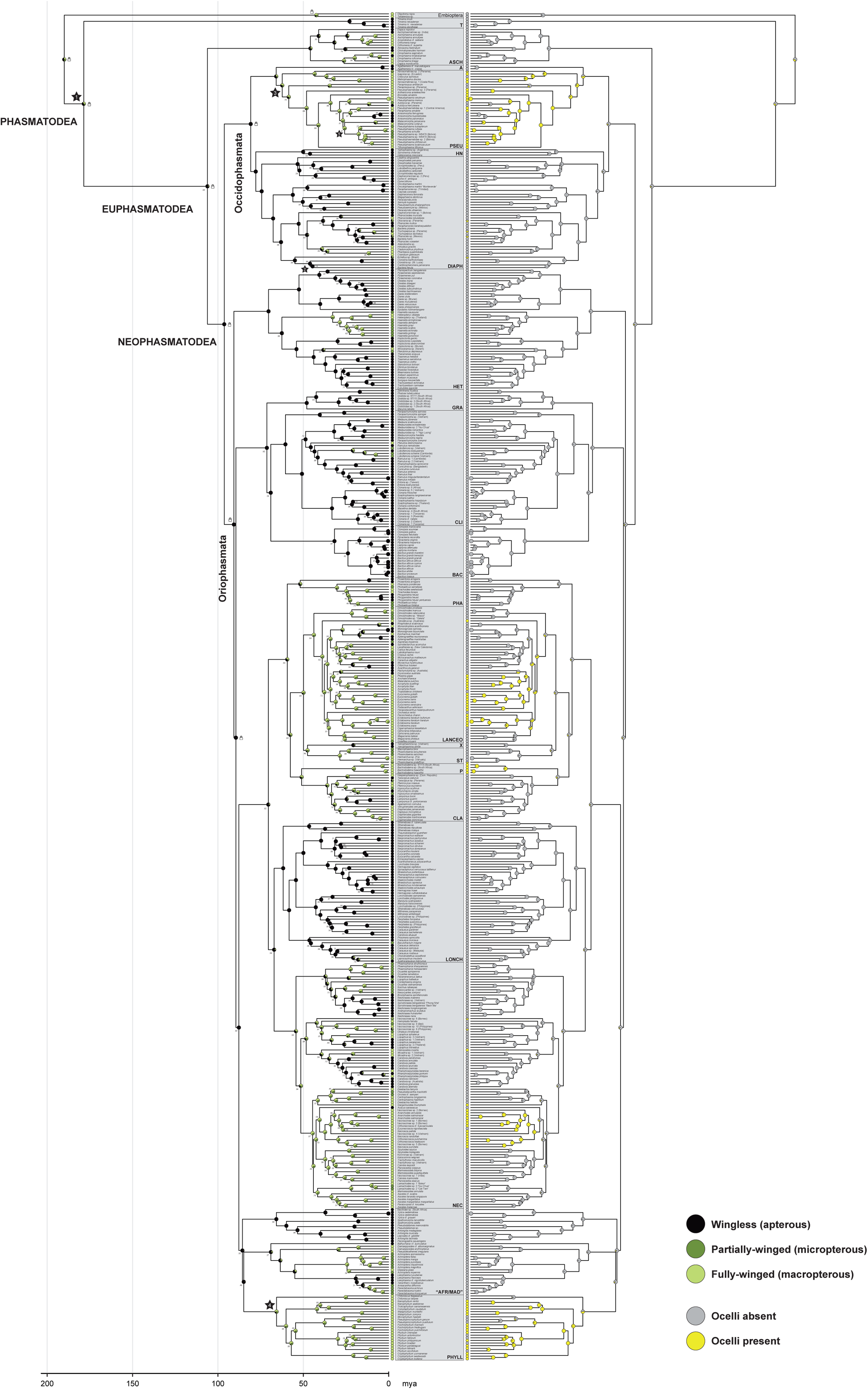
Ancestral state reconstruction for males based on the BI tree with B2 constraints (see lock symbols at nodes; Supplementary file 4). Nodal support values (<1 posterior probability) depicted at each node. Stars represent the fossils used for calibration and numbering corresponds to Supplementary file 16.

**Figure supplement 5.**
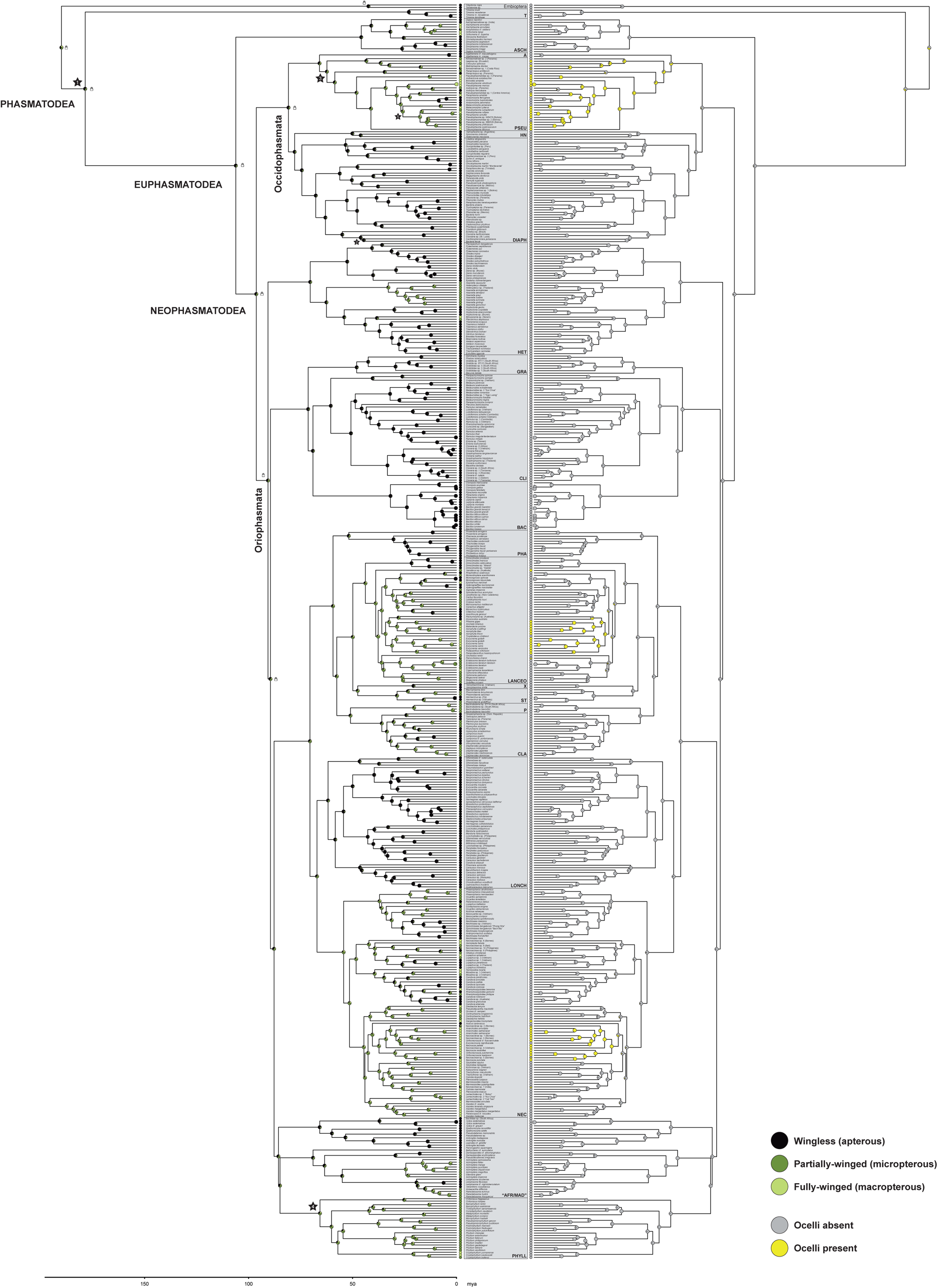
Ancestral state reconstruction for females based on the BI tree with B2 constraints (see lock symbols at nodes; Supplementary file 4). Nodal support values and divergence times are identical to those in Figure 4—figure supplement 4. Stars represent the fossils used for calibration and numbering corresponds to Supplementary file 16.

**Figure supplement 6.**
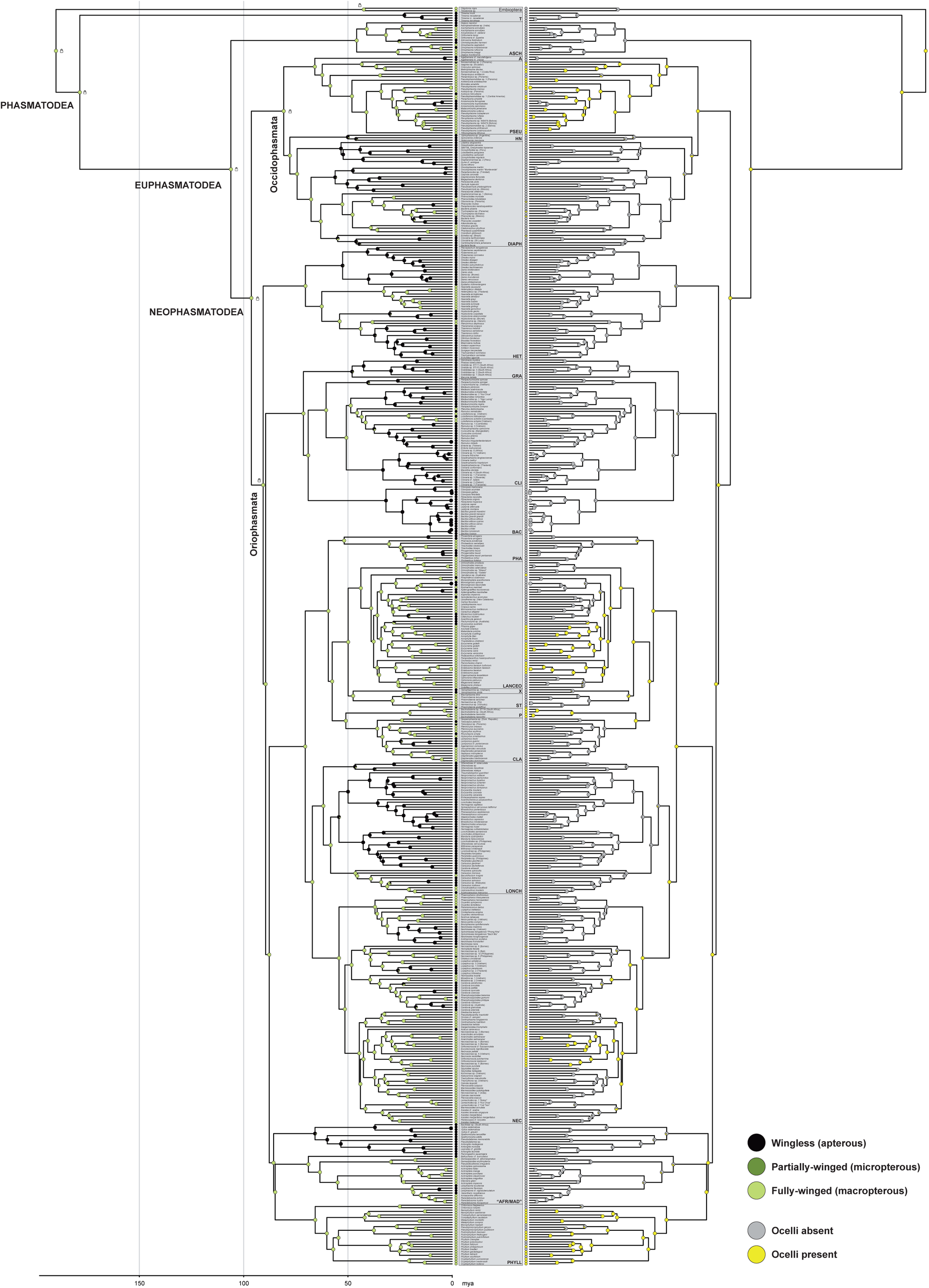
Ancestral state reconstruction for the binary dataset of males using the IRR model based on the BI tree with B2 constraints (see lock symbols at nodes; Supplementary file 4). Nodal support values and divergence times are identical to those in Figure 4—figure supplement 4. Stars represent the fossils used for calibration, and numbering corresponds to Supplementary file 16.

> **Source Data 1**. Results of stochastic character mapping. Numbers of transitions for ARD and IRR models. For the ARD model, the number of transitions from wingless to winged and vice versa are listed in addition to the combined number of transitions. The IRR is unidirectional, hence, the only transitions are from winged to wingless.

## Supplementary files

Supplementary file 1. Best scoring ML tree with UFBoot support values in newick format (TRE).

Supplementary file 2. Alternative ML tree with SH-aLRT support values in newick format (TRE).

Supplementary file 3. BI tree with B1 constraints in nexus format (NEX).

Supplementary file 4. BI tree with B2 constraints in nexus format (NEX).

Supplementary file 5. BI tree with B3 constraints in nexus format (NEX).

Supplementary file 6. Results of phylogenetic signal estimations using D statistics and Pagel’s lambda, and the comparison of the observed number of evolutionary transitions for each trait against a randomised character matrix. Wings = presence of wings (macropterous or micropterous), Flight = potential capability of flight (macropterous taxa), Ocelli = presence of ocelli. (XLSX).

Supplementary file 7. Results of testing the fit of different character evolution models to the binary wing dataset (data_wings_). ER, equal rates; ARD, all-rates-different; IRR, irreversible model disallowing the regain of wings. (XLSX).

Supplementary file 8. Probabilites [%] for the three wing states for the major nodes in B1, B2 and B3 constrained phylogenies. Note that Oriophasmata were not recovered as monophyletic in the B3 phylogeny and thus both Oriophasmata and Occidophasmata were excluded here. (XLSX).

Supplementary file 9. Average number of transitions between states resulting from 300 iterations of stochastic character mapping. Based on phylogeny with B2 constraints and both 3-states and binary wing and ocelli datasets using the ARD and the IRR model. The asterisk indicates that depicted ranges show the confidence interval of 95% highest posterior density. apt, apterous; micro, micropterous; macro, macropterous. (XLSX).

Supplementary file 10. Results of fitting Pagel’s model to test for correlation between the evolution of wings and ocelli. Correlation was tested on the binary datasets of males and females with different settings for the root state. Variable x represents the wing state and y the ocelli state. For males, the correlation of ocelli was further tested for the three wing states. Significance in bold. AIC, Akaike Information criterion, AICw, Akaike weights. (XLSX).

Supplementary file 11. Phylorate plots resulting from diversification rate estimation in BAMM. (A) Credible set of shift configurations with posterior probabilities (pp). Mean rate parameters are model-averaged across all samples assignable to a given configuration. (B) Phylorate plot of net diversification. Model shifts are depicted as symbols on branches, with star (=Euphasmatodea) and circle (=European Bacillinae) according to the best shift configuration. The red colouration indicates rate acceleration. The gray square represents an additional potential rate shift added from the second best configuration (clade includes Pharnaciinae+ *Prosentoria*, Palophidae, Cladomorphinae, *Xenophasmina*, Stephanacridini and Lanceocercata). (PDF).

Supplementary file 12. Results of fitting 24 models of diversification including HiSSE, BiSSE and character-independent (CID) models. Analysis was based on the B2 constraint phylogeny and the binary wing dataset with outgroup taxa (Embioptera) removed. Best-fit model is depicted in bold. AIC, Akaike information criterion. (XLSX).

Supplementary file 13. Information on taxon and gene sampling with GenBank accession numbers for each specimen used in this study. For taxa retrieved from GenBank, an ID with consecutive numbering is used (GBXXX). In alphabetical order. (XLSX).

Supplementary file 14. Final concatenated supermatrix in fasta format (FASTA).

Supplementary file 15. Best partitioning scheme (subsets) and best-fit substitution models determined with ModelFinder in IQ-TREE. (XLSX).

Supplementary file 16. Fossil calibrations used for the divergence time estimation in BEAST2. The numbering (1–5) corresponds to the depiction of calibrated nodes in Figure 4—figure supplement 2–5. (XLSX).

Supplementary file 17. Character matrix for females and males for ocelli and the 3-state and binary datasets for wings. Ocelli are coded as either absent or present and wings are coded in three states; apterous (0), micropterous (1) and macropterous (2). Missing data is shown as question mark. The binary datasets represent male wing states only and code presence of wings (data_wings_: wingless (0) vs. winged (1)) and flight capability (data_flight_: flightless (0) vs. flighted (1)). Missing data for binary datasets is coded as absent (0). For taxa retrieved from GenBank, an ID with consecutive numbering is used (GBXXX). (XLSX).

## Appendix

### Appendix 1. Phasmatodean phylogenetic relationships

#### Appendix 1A. Results

Our comprehensive taxon sample of 513 phasmatodean taxa and two outgroup species of Embioptera provided an optimal basis for exploring the phylogenetic relationships of Phasmatodea. All Maximum Likelihood (ML) and Bayesian (BI) analyses recovered highly congruent topologies with moderate to high support across the backbone nodes and for all the major lineages (Figure 4—figure supplement 1–4, Supplementary files 1–5). The application of different constraints had minimal effect on the topologies, and discrepancies largely pertained to the same groups such as the European Bacillinae and the African/Malagasy clade. When there was no constraints within Neophasmatodea (B3, Figure 4—figure supplement 3), the leaf insects (Phylliidae) were recovered as sister group to a clade of Occidophasmata + the remaining Oriophasmata, whereas the other inferences recovered them as closely related to the African/Malagasy group. Generally, the Occidophasmata are split in Agathemeridae + Pseudophasmatidae and Heteronemiinae + Diapheromerinae, and only in the ML tree *Paraprisopus* is recovered as sister taxon to *Agathemera* and not clustered within Pseudophasmatidae (Figure 4—figure supplement 1). The phylogenetic relationships within Oriophasmata vary slightly depending on the constraints. The most strict constraints forced the Heteropterygidae to form the sole sister group to the remaining Oriophasmata (B2, Figure 4— figure supplement 4), albeit the analyses without this constraint recovered them as sister taxon to a clade including Clitumninae *sensu* Cliquennois^1^ (= Clitumnini, Gratidiini, Medaurini), European Bacillinae *sensu* Cliquennois^1^ and African Gratidiidae *sensu* Cliquennois^1^. Yet, the B2 constraint resulted in the overall highest nodal support values. The remaining clade of Oriophasmata appears generally more congruent, especially in regard to the Lonchodidae (Lonchodinae + Necrosciinae) as sister to (Pharnaciinae + *Prosentoria*) + ((Palophidae + Cladomorphinae) + (Stephanacridini + *Xenophasmina* + Lanceocercata)).

The selection of fossils to use as calibration age priors was identical for all three BI trees (Supplementary file 16) and the resulting divergence time estimates are consistent among the analyses, with slightly older estimates for the B2 constrained tree (Supplementary files 3–5). Our results estimate Phasmatodea to have originated in the Jurassic ∼178.56 million years ago (mya) (186.6–165.14 mya) (B2), whereas the diversification of Euphasmatodea and Neophasmatodea started in the Cretaceous ∼106.13 mya (114.16–75.11 mya) and ∼96.29 mya (141.77–63.23 mya), respectively. However, the divergence of most of the major lineages does not predate the KP boundary and which thus have their origin in the Early Cenozoic (Palaeocene).

#### Appendix 1B. Discussion

While the phasmatodean taxonomy was long considered as problematic and unsolved especially in regard to other major insect lineages^2–5^, studies of recent years have illuminated much of the controversially discussed phylogenetic relationships^6–12^. Our study contributes to this progress by inferring a large-scale phylogenetic analysis of over 500 phasmatodean taxa including sequence data for nearly 200 so far unsequenced species. Despite the limited number of genes in comparison to phylogenomics studies, our results show promising similarities in the overall topology.

The New World clade Occidophasmata that was only recently revealed by phylotranscriptomic studies^7,8^, was highly supported by our phylogenetic inferences, even when no constraints were set (B3 constraint; Figure 4—figure supplement 3), with *Agathemera* as sister taxon to Pseudophasmatidae^7,8,13^. Our increased taxon sampling furthermore revealed the Heteronemiinae, which have not yet been included in any phylogenomic study, as closely related to Diapheromerinae^14^ and not as previously thought to Pseudophasmatidae^1,15,16^. By contrast, our inferences could not resolve the similarly contradicting phylogenetic position of Paraprisopodini proposed to be a clade of or closely related to Pseudophasmatidae^6,16–18^, Diapheromerinae^19^ or unrelated to these neotropical lineages^20^. Our ML inference recovered *Paraprisopus* as sister taxon to *Agathemera* (Figure 4—figure supplement 1), while their position within Pseudophasmatidae in the BI trees (Figure 4—figure supplement 2–4) is biased due to the constraint set on Pseudophasmatidae (incl. *Paraprisopus*) for the fossil calibration.

Further studies including additional members of Paraprisopodini as well as Prisopodini are needed to examine the phylogenetic relationships. That a more extensive review of the Neotropical taxa is needed was already demonstrated by Robertson et al.^6^, who transferred *Otocrania* formerly assigned to Cladomorphinae to the Diapheromerinae. Bank et al.^10^ found the nominal taxon *Cladomorphus* to cluster among Diapheromerinae as well, and here we additionally recover two cladomorphine members (*Cranidium* and *Hirtuleius*) within Diapheromerinae suggesting that the members of Cladomorphinae *sensu stricto*^21^ in fact belong to the Diapheromerinae or Bacteriidae *sensu* Cliquennois^1^. Consequently, the remaining Cladomorphinae (Haplopodini, Hesperophasmatini, Pterinoxylini)^21^ were suggested to be named Haplopodidae^1^. Furthermore, we show that also the Brazilian *Echetlus* appears to be a member of Diapheromerinae as proposed in earlier morphological studies^22,23^ indicating that this taxon is not congeneric with the Southeast Asian members of *Echetlus* or the Australian Necrosciinae *Candovia* to which some of its members were later assigned^24^.

Among the Old World Oriophasmata, we found much congruence with previous molecular phylogenies regarding the phylogenetic relationships of Lonchodidae (Necrosciinae + Lonchodinae)^6^ and their sister taxon comprising Palophidae, Cladomorphinae (excl. *Cladomorphus, Cranidium* and *Hirtuleius*; see above), Pharnaciinae, Stephanacridini, *Xenophasmina* and Lanceocercata^6–8,10,11^. Within the latter, our results support the recently established Megacraniinae^22^ (excl. *Apterograeffea*), albeit assuming a subordinate position as opposed to the proposed relationships by Hennemann^22^. Besides the well-supported sister groups from the Mascarene Islands (Monandropterinae *sensu* Cliquennois^1^) and from New Zealand and New Caledonia (Acanthoxylinae *sensu* Cliquennois^1^), the remaining clades of Lanceocercata (i.e., Phasmatinae, Pachymorphinae, Tropidoderinae and Xeroderinae *sensu* Cliquennois^1^) appear as highly polyphyletic^6,9,14,25,26^ and are in need of formal taxonomic revision. This is particularly true with regard to the Xeroderinae, which have repeatedly been shown to be polyphyletic, and whose member *Xenophasmina* we recovered to either form the sister group to the remaining Lanceocercata (Figure 4—figure supplement 1 and 3) or to be entirely unrelated to Lanceocercata and closer related to Stephanacridini (Figure 4—figure supplement 2 and 4)^7^.

The phylogenetic relationships of the remaining lineages of Oriophasmata appear rather inconsistent with the results of previous molecular analyses^6–8^. For instance the Clitumninae *sensu* Cliquennois^1^ (Clitumnini + Gratidiini + Medaurini) were repeatedly shown as closely related to Pharnaciinae^7,8,13^, but were here recovered in close relationship with the European Bacillinae and the African Gratidiidae *sensu* Cliquennois^1^, and with the Heteropterygidae, when no constraints were set (Figure 4—figure supplement 1–3). The low support for their relatedness, especially when Heteropterygidae are included, suggests that the topology might not be reliable. Since most of the bacilline taxa included in our inferences are represented by only one or two genes, the inferred phylogenetic position of this European group of stick insects is potentially biased, and a closer relationship to the Malagasy clade is favoured as is corroborated by phylogenomic studies and in terms of biogeography^7,8^. By contrast, the Gratidiidae (or any African taxon in general) have not yet been included in a phylogenomic study and their placement among Oriophasmata remains unclear, though Robertson et al.^6^ recovered members of this lineage as close relatives to the Malagasy stick insects. In our inferences, however, the African Xylicinae are revealed as closely related to the Malagasy group. In fact, *Xylica* and an unidentified Bacillidae (possibly Xylicinae) species are highly supported as sister group to the Antongiliidae + *Spathomorpha*, while the xylicine *Bathycharax* appears unrelated to this assemblage, but is recovered as sister taxon to either the remaining Malagasy stick insects (Figure 4—figure supplement 2 and 4) or to the whole Africa/Madagascar clade (Figure 4— figure supplement 1 and 3). The African taxa are generally highly underrepresented in molecular analyses and the inclusion of several species in our study did not succeed in illuminating their evolutionary history with exception of the Palophidae, which were already previously and repeatedly recovered as sister taxon to the Cladomorphinae^6,10,11,14^. The taxonomic shortcomings may be overcome by future work focussing on the underrepresented taxa of Africa and its associated regions. A more comprehensive taxon sampling will also be needed to elucidate the historical biogeography revolving around the colonisation of Madagascar, in particular, since our results are inconclusive about the monophyly of an African/Malagasy lineage with respect to the leaf-imitating Phylliidae. Although our results partly support the hypothesis that phylliids are closely related to European, African and Malagasy stick insects as was proposed by the phylotranscriptomic results by Simon et al.^7^ in accordance with the ancestral distribution of leaf insects in Europe^27^, clarification is needed to fully understand the life history of these lineages including the origin and the controversially discussed sister group of leaf insects^7,8,11^.

**Appendix-figure 1.**
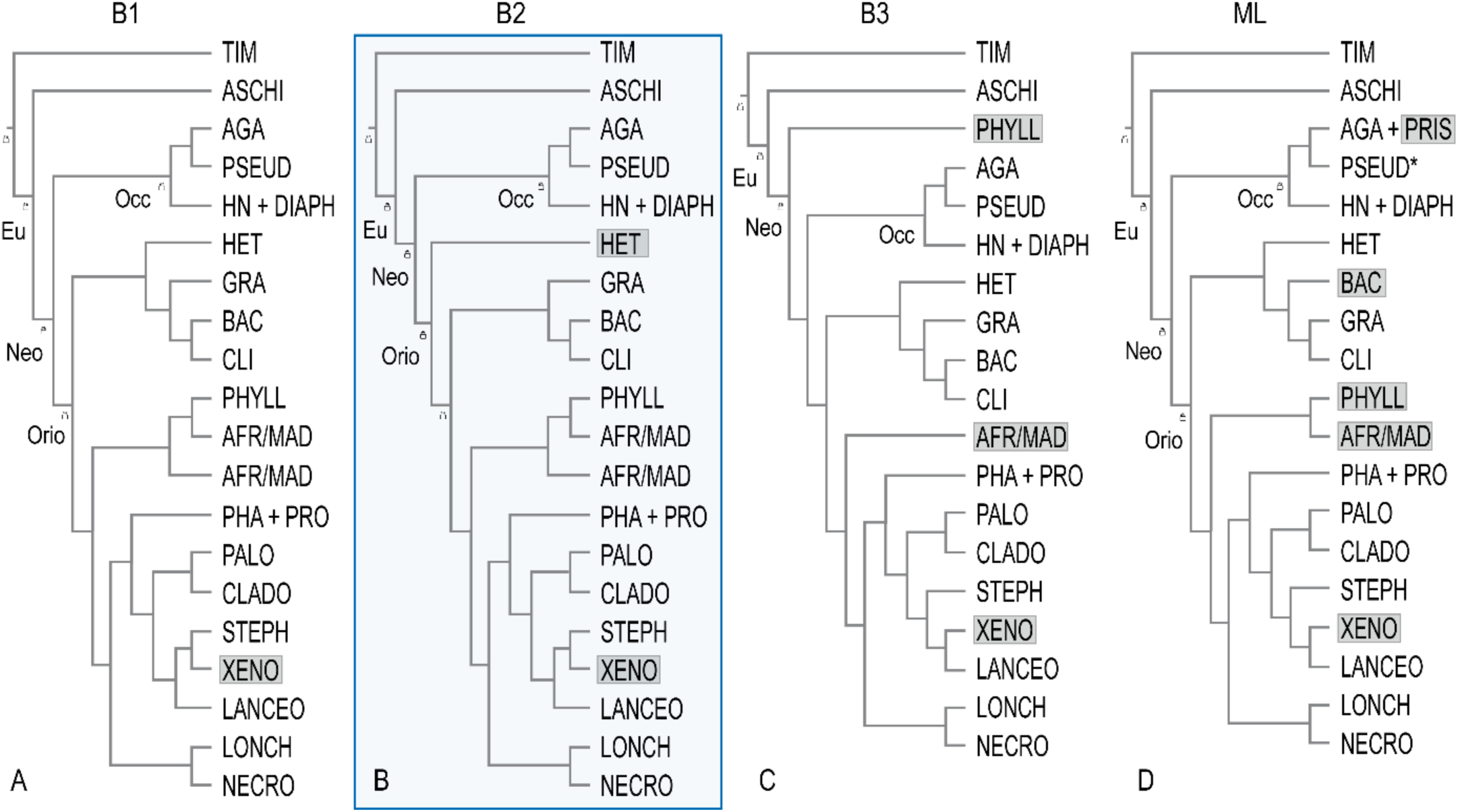
Comparison of the four inferred phylogenies based on ML and BI methods as well as different topological constraints (B1–B3) depicted as lock symbols at the nodes (Figure 4—figure supplement 1–4). Taxa holding a different position in comparison to the other topologies are highlighted. Blue frame indicates the topology on which subsequent analyses are based. Eu, Euphasmatodea; Neo, Neophasmatodea; Orio, Oriophasmata; Occ, Occidophasmata; TIM, Timematodea; ASCHI, Aschiphasmatidae; AGA, Agathemeridae; PSEUD, Pseudophasmatidae including *Paraprisopus*; PSEUD*, Pseudophasmatidae excluding *Paraprisopus*; HN, Heteronemiinae; DIAPH, Diapheromerinae; HET, Heteropterygidae; GRA, Gratidiidae *sensu* Cliquennois (2020); BAC, Bacillinae *sensu* Cliquennois (2020); CLI, Clitumninae *sensu* Cliquennois (2020); AFR/MAD, African/Malagasy group including Achriopteridae, Anisacanthidae, Antongiliidae, Damasippoididae and Xylicinae *sensu* Cliquennois (2020); PHYLL, Phylliidae; PHA, Pharnaciinae + *Prosentoria*; LANCEO, Lanceocercata; XENO, *Xenophasmina*; STEPH, Stephanacridiini; PALO, Palophidae; CLADO, Cladomorphinae; LONCH, Lonchodinae, NECRO, Necrosciinae.

